# Evaluating the performance of protein structure prediction in detecting structural changes of pathogenic nonsynonymous single nucleotide variants

**DOI:** 10.1101/2023.11.25.568523

**Authors:** Hong-Sheng Lai, Chien-Yu Chen

## Abstract

Protein structure prediction serves as an efficient tool, saving time and circumventing the need for laborious experimental endeavors. Distinguished methodologies, including AlphaFold, RoseTTAFold, and ESMFold, have proven their precision through rigorous evaluation based on the last Critical Assessment of Protein Structure Prediction (CASP14). The success of protein structure prediction raises the following question: can the prediction tools discern structural alterations resulting from single amino acid changes? In this regard, the objective of this study is to assess the performance of existing structure prediction tools on mutated sequences. In this study, we posited that a specific fraction of the pathogenic nonsynonymous single nucleotide variants (nsSNVs) would experience structural alterations following amino acid mutations. We meticulously assembled an extensive dataset by initially sourcing data from ClinVar and subsequently applying multiple filters, resulting in 964 alternative sequences and their corresponding reference sequences. Utilizing UniProt, we acquired reference sequences and generated the corresponding alternative sequences based on variant information. This study performed three tools of structure prediction on both the reference and alternative sequences and expected some structural changes upon mutations. Our findings affirm AlphaFold as the foremost prediction tool presently; nonetheless, our experimental results underscore persistent challenges in accurately predicting structural alterations induced by nsSNVs. Discrepancies between the predicted structures of reference and alternative sequences, when observed, often stem from a lack of confidence in the predictions or the spatial separation between compact domains interrupted by disordered regions, posing challenges to successful alignment.

## 1 Introduction

Protein structure prediction is the endeavor to anticipate the three-dimensional structure of a protein based on its amino acid sequence. This pursuit holds paramount importance in bioinformatics and genomics, bearing substantial implications for medical applications such as drug design [1]. This approach has substantially mitigated the time and labor-intensive nature of deciphering protein structures, traditionally achieved through X-ray crystallography, cryo-electron microscopy structure modeling, and solution nuclear magnetic resonance spectroscopy [2]. In the current era dominated by deep learning, there has been a noteworthy enhancement in prediction accuracy. An increasing array of models has been deployed in real-world studies, marking a significant advancement in the field.

Every two years, the performance of protein structure prediction tools is evaluated through Critical Assessment of Protein Structure Prediction (CASP) [3], which serves as a critical benchmark for protein structure prediction techniques. In 2020, a ground-breaking protein structure prediction tool, AlphaFold2 [4], developed by the Google DeepMind team, achieved remarkable success, obtaining a score of 92.4 out of 100 in CASP14, a substantial leap from the previous accuracy levels of around 40 out of 100. Similarly, within the same year, the RoseTTAFold [5] tool developed by David Baker’s team from the University of Washington achieved lower yet comparable predictive performance using a smaller dataset and faster prediction times. Both tools employ multiple sequence alignment (MSA) to archive homologous sequences in databases for reference. Following identifying similar sequences, an attention model is employed to predict the three-dimensional protein structure, followed by refinement based on the atomic chemical properties. Moreover, in 2022, ESMFold [6], developed by Meta AI, demonstrated the ability to directly infer the protein structure from the primary sequence using a large language model (LLM) with up to 15 billion parameters. ESMFold accelerated at least six times more than AlphaFold2 in the AI inference phase, enabling the construction of a large-scale metagenomics protein data bank. All the generated protein structures from the previously mentioned protein structure prediction tools claim high similarity to actual protein structures, achieving accuracy at the atomic level, thereby aiding in the further determination of the actual function of the protein structure.

When a DNA sequence changes a single nucleotide—represented by the nucleotides A, T, C, or G—a single nucleotide variant (SNV) emerges, constituting the most prevalent type of sequence variation. Among SNVs, synonymous changes maintain the amino acid sequence unaltered, whereas nonsynonymous single nucleotide variants (nsSNVs) introduce modifications to the amino acid sequence, consequently impacting the protein’s functionality [7]. However, elucidating the influence of nsSNVs on protein function proves challenging in clinical studies [8]. Even being annotated as pathogenic variants, the actual impact of nsSNVs on protein folding, binding, expression, and other protein features remains uncertain and necessitates further investigation [9].

In contemporary research, the predominant focus has been predicting the pathogenicity or thermodynamic free energy of nsSNVs rather than their structural changes [10]. Prior to the emergence of AlphaFold2, the confidence in protein structure prediction results was insufficient. So, when predictive data was available, it held limited value for further discussions [11]. With the advent of various high-precision prediction models, some studies employed protein structure prediction tools to investigate nsSNVs from only under 30 genes for analysis [12]. On the other hand, some studies asserted the inability to predict non-wildtype sequences through structural prediction tools, yet lacking substantial evidence and experiments about the claim [13]. We aspire to expand the scope of prediction to encompass pathogenic nsSNVs in humans using protein structure prediction tools. This endeavor is expected to contribute significantly to structural biology and medicine, facilitating the deduction of their pathogenic mechanisms.

In this study, we compiled a comprehensive pathogenic nsSNVs dataset. With the hypothesis that a specific fraction of the pathogenic nsSNVs would experience structural alterations following amino acid mutations, we expected to observe some structural changes on mutated sequences against the reference sequences. Three tools are employed in this study: AlphaFold, RoseTTAFold, and ESMFold. Through the profound impact of protein structure prediction tools on the scientific community, this study aspires to apply these tools to the context of pathogenic variants, aiming to enhance our understanding of the functional consequences of nsSNVs.

## 2 Methods

### 2.1 Dataset

We selected nsSNVs from the ClinVar [14], an open and accessible repository containing records detailing the connections between human genetic variations and observed health status. The classification of pathogenicity includes five categories: Pathogenic, likely pathogenic, uncertain, likely benign, and benign. We have chosen to focus on the “Pathogenic” category for discussion and ensure the nsSNVs are from multiple submitters. In this step, we selected 4,281 variants (Figure 1).

**Figure 1:**
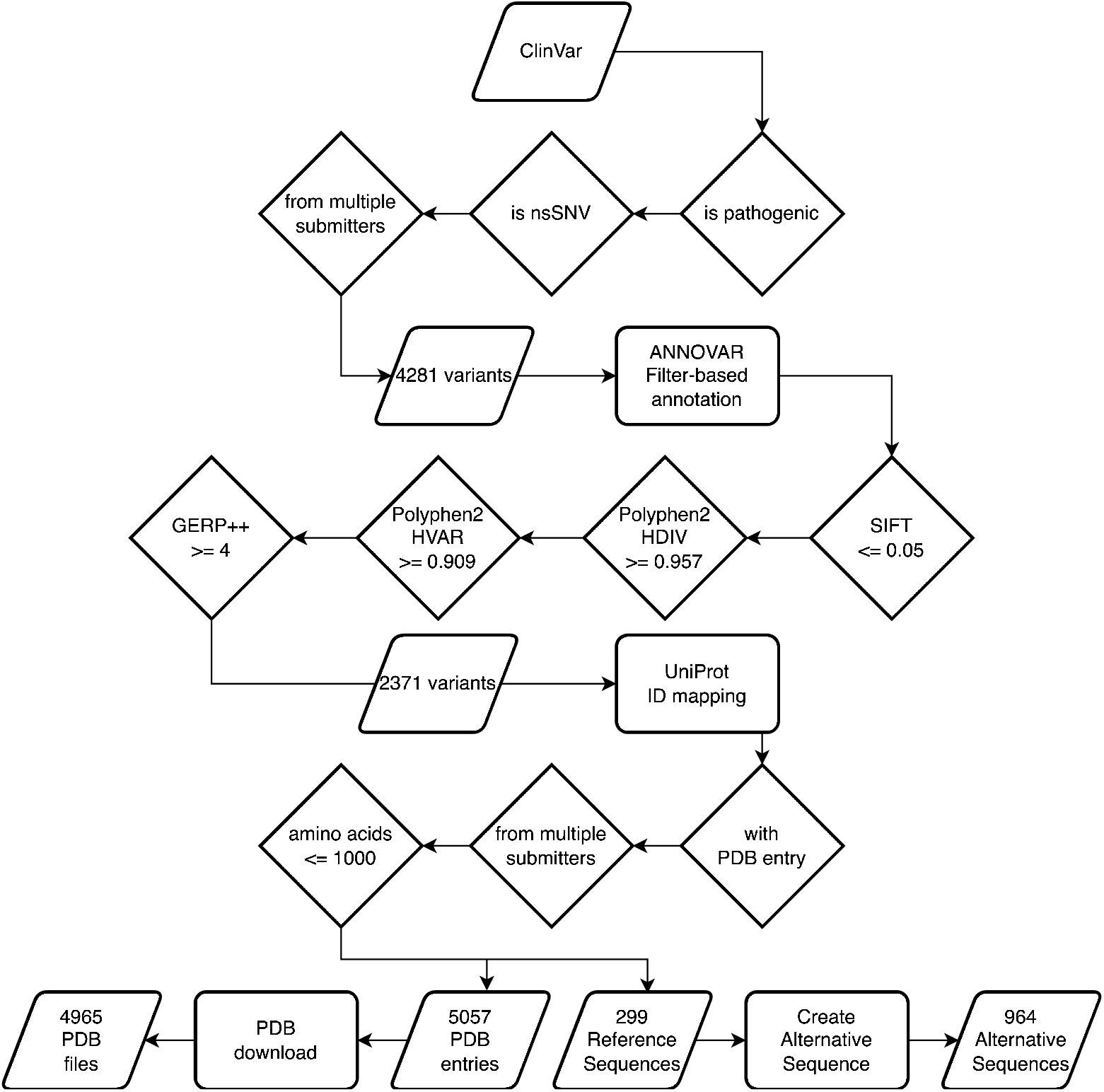
Flow chart of obtaining pathogenic nsSNVs, experimental structures in PDB format, and reference sequences.

Subsequently, we applied a filtered-based annotation database in ANNOVAR [15] to functionally annotate genetic variants. Two kinds of filtering were undertaken to ensure the variants’ impact on structural changes and the conservation of these variants. To select the variants with possible impact on structural changes, we employed SIFT [16], Polyphen2_HDIV, and Polyphen2_HVAR [17]. On the other hand, to retain high conservation variants, we adopted GERP++ score [18]. For Polyphen2, the HDIV score was selected to assess rare alleles at potentially implicated loci in complex phenotypes, dense mapping of regions identified by genome-wide association studies, and the analysis of natural selection using sequence data. Variants with a score >= 0.957 were considered. Additionally, we employed the HVAR score, which aims to distinguish mutations with significant effects from the broader spectrum of human variation, encompassing mildly deleterious alleles. Variants with a score >= 0.909 were chosen. Regarding SIFT, we utilized a threshold of score <= 0.05 to identify nsSNVs predicted to be deleterious. All selection criteria were derived from thresholds provided in the relevant literature, and these scores were tied to evaluations of protein structural impact on determining deleteriousness. We understand that Polyphen2 and SIFT scores are not among the top-performing indicators in pathogenicity score prediction now. However, these scores have a more direct relationship with structural variations in their training stage than other top-performing ensemble models. Next, variants with GERP++ scores >= 4 are kept. Generally, the higher the score, the more conserved the site, while the overall scores range from −12.3 to 6.17. Eventually, we obtained 2,371 variants characterized by high conservation and a strong correlation with structural variations (Figure 1).

After these filtering steps, we identified reference sequences in Universal Protein Knowledgebase (UniProt) [19] for the selected variants and opted for source files with contributions from multiple submitters. To enable further comparisons, we retained reference sequences with corresponding experimentally determined structure entries for the Protein Data Bank (PDB) [20]. Considering computational efficiency, we chose sequences shorter than 1,000 amino acids as the final dataset. We eventually found 299 reference sequences and created 967 alternative sequences based on the variant details (Figure 1). The same reference sequence may correspond to different variant sequences of nucleic acids. Three of the 967 variant sequences are attributed to start-loss mutations. Due to the uncertainty regarding the meaning of their sequences, these particular mutations were excluded. Additionally, certain entries corresponding to PDB were not considered due to their unavailability for download. Eventually, we have 964 alternative sequences for the experiment.

### 2.2 Multiple Sequence Alignment for Experiment Data

For each reference sequence, multiple corresponding PDB entries might exist. To determine which entries are the most similar to the reference protein sequences, we used ClustalW [21] to perform MSA. Firstly, we used DBREF records from the PDB files to identify whether the reference was from the same UniProt reference. We then extracted the SEQRES records from these files, representing the sequences researchers intended to observe through experimental methods. We also extracted the ATOM records from the files, representing the actual observed protein structures, and converted them into sequences for further comparison. We aimed to ensure only one reference structure source in the PDB file and a single protein structure. This assertion helped us guarantee that our protein structure alignment in the subsequent steps remained undisturbed by other factors. By comparing these extracted sequences with the wild-type sequences from UniProt, we identified the experimentally determined structures most closely aligned with the sequences we wanted to compare by similarity scores.

### 2.3 Protein Structure Prediction

Protein structure prediction has three stages: sequence representation generation, artificial intelligence inference, and protein structure relaxation (Figure 2). Among the three tools we are comparing AlphaFold, RoseTTAFold, and ESMFold the most significant divergence lies in sequence representation generation. AlphaFold utilizes MSA to generate homologous sequences, whereas RoseTTAFold employs cropped MSA, significantly reducing time but compromising accuracy. ESMFold, on the other hand, employs a 15-billion-parameter LLM as a pre-trained model. It transforms the amino acid sequence into a sequence representation vector, followed by AI inference and relaxation stages identical to AlphaFold.

**Figure 2:**
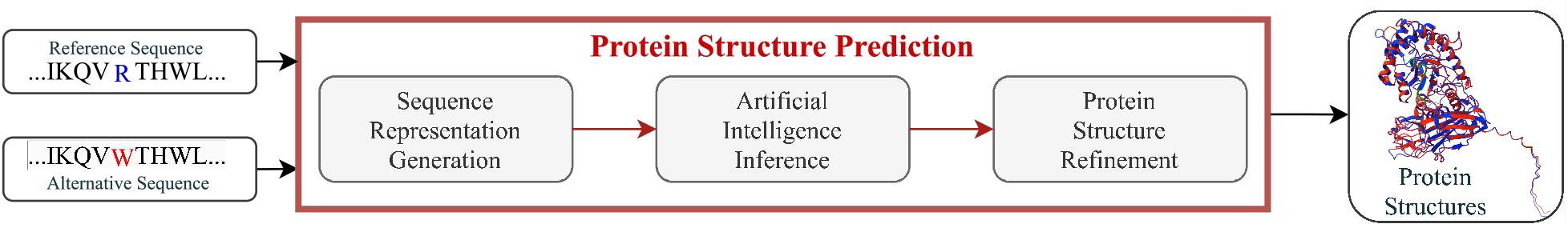
Protein Structure Prediction Pipeline. The blue amino acid represents the position in the reference sequence where the variation occurs, while the red amino acid indicates the mutated amino acid. The example of the predicted protein structures is plotted after structure alignment. The blue structure represents the result predicted based on the reference sequence, while the red structure represents the result predicted based on the variant sequence.

In this study, to assess the impact of MSA on AlphaFold, we also ran AlphaFold without utilizing MSA as input, considering only the reference sequence. We also investigated whether the MSA depth in AlphaFold affects the confidence region scores the pLDDT score. Herein, pLDDT >= 90 corresponds to high confidence, 90 > pLDDT >= 70 indicates confidence, 70 > pLDDT >= 50 implies low confidence, and pLDDT < 50 corresponds to very low confidence. Very low-confidence predictions are often associated with intrinsically disordered proteins.

We ran AlphaFold v2.3.1, RoseTTAFold v1.1.0, and ESMFold from ESM v2.0.0 in TWCC High-Performance Computing cloud service, using 1 Tesla V100 GPU (32GB VRAM), 6 CPU, 90 GB memory for each sequence-tostructure prediction.

### 2.4 Protein Structure Alignment

TM-align [22] is an algorithm employed for the optimal structural alignment of proteins. The outcomes of TM-align encompass three intuitive pieces of information: Template modeling score (TM-score) [23], Root Mean Square Difference (RMSD) [24], and alignment length. The primary assessment criterion in this study is TM-score, supplemented by RMSD.

The main drawback of RMSD is its susceptibility to strong fragment errors. Additionally, RMSD is influenced by the length of alignment (*L*_*aligned*_), making it an unsuitable metric for variable length or global protein sequence alignments, with the equation:

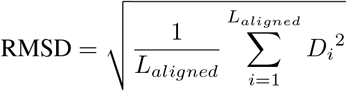

where *D*_*i*_ is the distance between the *i*-th pair of residues in the reference and alternative structures.

This issue might not be apparent when comparing predicted reference structures with alternative structures, as we can ensure that the lengths of the two structures are equal. However, when contrasting a reference structure with an experimental structure, we cannot guarantee the alignment of the lengths on both sides.

The TM-score aims to address the limitations of RMSD, perform global assessment analysis, and enable comparison among protein models of varying amino acid lengths. It can be told from the equation that this is a normalized formula based on experimental results. Due to the adjustments, the dependency on protein length represented by *L*_*target*_ is reduced, and the equation is:

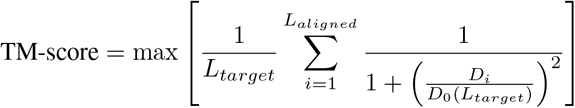

where 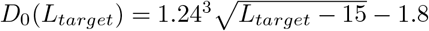 is a scaling factor determined by experiment.

Moreover, because of the globally evaluative nature of TM-score, it establishes a rational method for numerical comparison. This aspect is a crucial indicator in this research project. When a score is below 0.2, it is equivalent to aligning two randomly unrelated proteins, while a score exceeding 0.5 indicates a certain degree of similarity between the two structures with analogous three-dimensional folding arrangements. We utilized these two thresholds as the demarcation criteria for assessing alignment effectiveness in this study.

Normalization is conducted based on the experimental structure when comparing the experiment-generated structure. On the other hand, if the comparison involves a reference structure and an alternative structure, normalization is conducted with respect to the reference structure.

## 3 Results

### 3.1 Prediction Analysis

In Figure 3, we observed the execution time required for reference sequences by the three tools. Due to the fast processing time of ESMFold, it is not visible using second as the time scale. In this regard, the plot is represented using logarithmic time. Across all prediction tools, the time required follows the order Alphafold > RoseTTAFold > ESMFold in all cases. The absence of colored segments indicates instances where the prediction tool failed to determine the protein structure. ESMFold, given its utilization of sequence representation generated by language models, can predict structures as long as there is sufficient memory. Its practical limit is approximately 850 amino acids (in 32G VRAM devices). RoseTTAFold primarily encounters memory-related issues, either due to the maximum matching number constraints on the MSA or inadequate AI inference memory. In the original version of AlphaFold, structural prediction failures were not observed; instead, longer sequences resulted in exponential execution time without forced termination due to memory constraints. However, in the latest version (v2.3.1), early termination may occur during the sequence representation stage due to MSA tool-related issues. We also compared AlphaFold without MSA to the original AlphaFold. Figure 3(b) shows that the MSA version significantly reduces the time required, indicating that a substantial portion of the processing time is spent on MSA computation. Additionally, we observed from ESMFold and AlphaFold that without MSA, AI inference time is highly correlated with sequence length, the primary source of time uncertainty.

**Figure 3:**
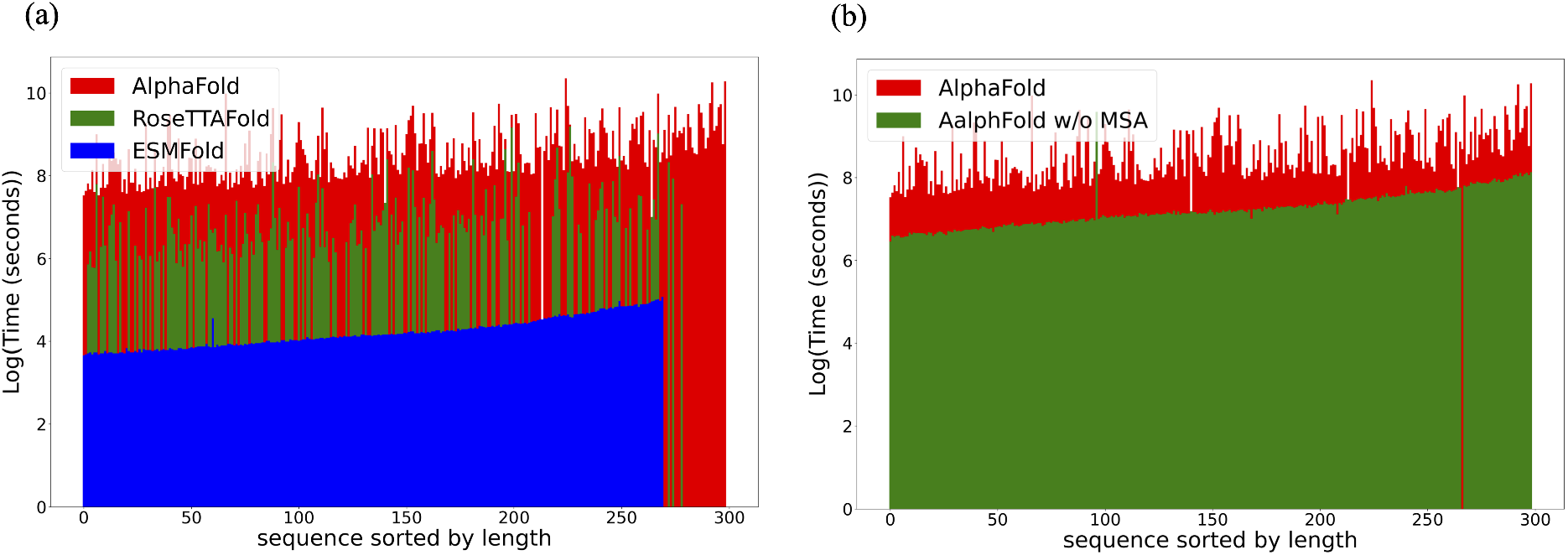
Time analysis for three protein structure prediction tools. (a) The disappeared bars in the plots represent cases where the prediction tool failed to predict successfully. Among them, ESMFold can complete the predictions until approximately 850 amino acids. (b) Time comparison between AlphaFold and AlphaFold without MSA. It can be observed that the sequences of the prediction failures for both methods are different, and the time required AlphaFold > AlphaFold without MSA in all cases.

Regarding AlphaFold, we aimed to delve into the details of MSA. We have observed a high similarity between the MSA of reference sequences and their corresponding variant sequences (Figure 4(a)). To be more precise, we found that in 403 instances, the MSA depth of reference sequences is smaller than the MSA depth of variant sequences. In contrast, in 543 cases, the MSA depth of variant sequences is smaller than that of their corresponding reference sequences. Additionally, there was one instance where the MSAs were entirely identical. Furthermore, it was noteworthy that all the MSAs with smaller depths were entirely contained within the larger MSAs. This indicated that single nucleotide variations had minimal impact on MSA, subsequently affecting downstream AI inference. As depicted in Figure 4(b), we also observed that the sequence length did not significantly influence the depth of MSA.

**Figure 4:**
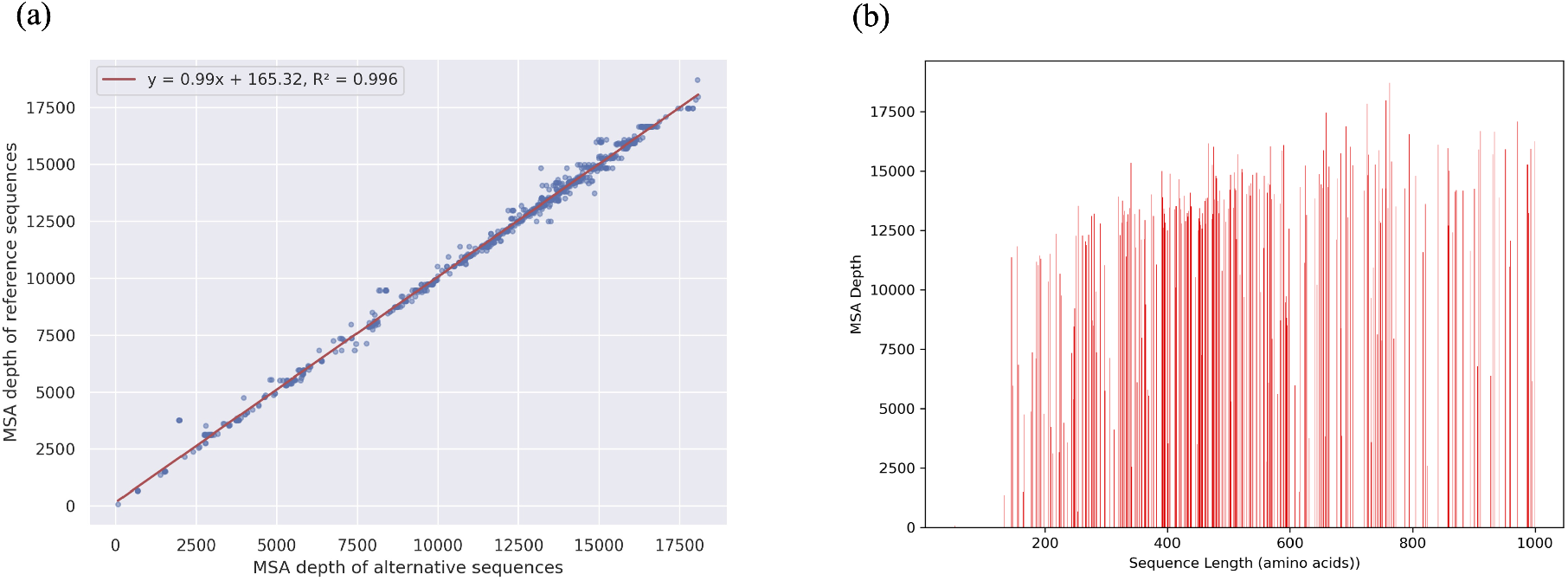
Number of sequences in AlphaFold MSA. (a) Correlation between AlphaFold MSA depth of an alternative sequence versus its corresponding reference sequence. (b) MSA depth for AlphaFold in reference sequences.

### 3.2 Predicted structures vs. Experimental structures

We provided the accuracy of the protein structure prediction tools in Table 1. First, we compared all PDB structures with the predicted structure of a reference sequence. It can be seen that all three tools are able to predict protein structures with high precision, as claimed. When we selected the PDB structure with the most similar sequence after performing ClustalW, it was evident that the average results showed an improvement. The detailed distribution of the TM-score can be observed in Table 2. Even though RoseTTAFold seems to have the best overall performance, it infers much fewer predicted structures. On the other hand, ESMFold, despite having the worst performance in TM-score, yields a higher number of successful predictions. Overall, AlphaFold remains one of the most accurate tools currently available after we compared the intersection of ‘Best PDBs’ sets of the three tools, and thus, it is the primary tool for the following focused discussion.

**Table 1:** TM-scores between predicted structures of reference sequences and experimental structures in PDB. (The number in parentheses represents the number of structures to evaluate.)

| Average TM-score | AlphaFold | AlphaFold w/o MSA | RoseTTAFold | ESMFold |
| --- | --- | --- | --- | --- |
| All PDBs (total 4965) | 0.8069 (4933) | 0.7983 (4960) | 0.8184 (3286) | 0.7923 (4580) |
| Best PDBs (total 159) | 0.8352 (157) | 0.7983 (158) | 0.8359 (78) | 0.8068 (141) |
| Intersection of Best PDBs | 0.8791 (73) | 0.8403 (73) | 0.8426 (73) | 0.8626 (73) |

**Table 2:** Number of samples in different TM-score ranges between predicted structures of reference sequences and best experimental structures in PDB.

| TM-score (t) | $t < 0.5$ | $0.5 \leq t < 0.7$ | $0.7 \leq t < 0.9$ | $0.9 \leq t$ | Total |
| --- | --- | --- | --- | --- | --- |
| AlphaFold | 15 | 16 | 42 | 84 | 157 |
| AlphaFold w/o MSA | 23 | 21 | 32 | 82 | 158 |
| RoseTTAFold | 6 | 8 | 19 | 45 | 78 |
| ESMFold | 15 | 16 | 41 | 69 | 141 |

From the corresponding numbers for AlphaFold without MSA, we can deduce that a portion of AlphaFold’s failures to infer structures successfully is in the MSA stage. Even though the numerical values are the same, the average TM-score still slightly improved, indicating that our method of selecting the best structures for comparison is beneficial in the evaluation. It’s also evident that completely omitting MSA impacts the model’s performance because the AlphaFold can only rely on template structures. Another evaluative aspect is the model’s confidence level in its own structures, as represented by the pLDDT score. In terms of average pLDDT scores, the ranking is as follows: AlphaFold (83.25) > AlphaFold without MSA (80.17) > ESMFold (79.27) > RoseTTAFold (72.73). It is still evident that AlphaFold has a higher confidence level in its predicted structures.

When we compared protein structure prediction tools against all experimental structures with TM-score less than 0.5, it became apparent that most poorly predicted structures exhibit significant overlap (Figure 5(a)), showing some of the structures still cannot be predicted by currently existing tools. From Figure 5(b), we observed that the proportion of aligned lengths is never greater than the TM-score owing to the definition. Furthermore, the distribution trend indicated that the higher the aligned length, the higher the corresponding TM-score.

**Figure 5:**
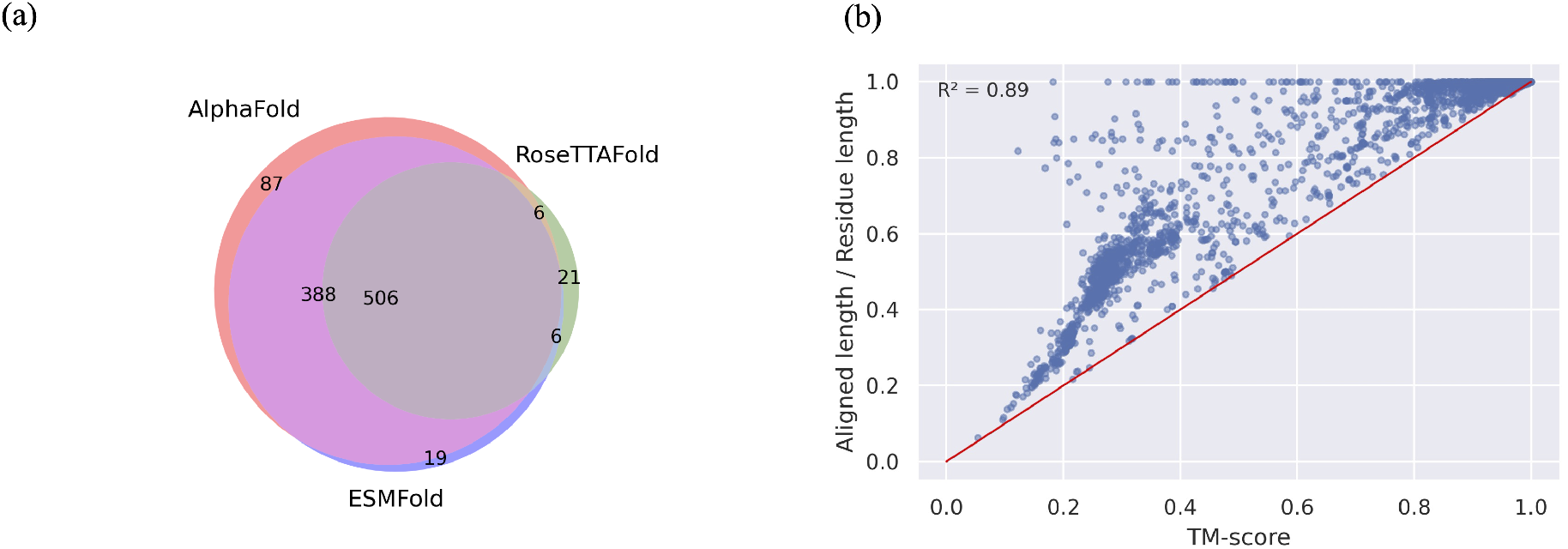
Poor predictions in three protein structure prediction tools. (a) A Venn diagram illustrating significant overlap of poor predictions (TM-score < 0.5 when compared to experimentally determined protein structures) among protein structure prediction tools. (b) Correlation between TM-score and the proportion of aligned length (aligned length / total residue length) in AlphaFold predicted reference structures and experimentally determined protein structures. The red line’s slope is 1.

### 3.3 Prediction of reference sequences vs. alternative sequences

Next, we further discuss the structural similarity between the predicted structures of reference sequences and the predicted structures of alternative sequences (Figure 6). It can be seen that most of the predicted structures remain highly similar (TM-score > 0.9) between reference and alternative. Among the three tools, ESMFold exhibits the most significant similarity. This is because it relies solely on natural language processing techniques and lacks any variations derived from MSA, resulting in less input variability than the other tools, which provide little assistance for subsequent AI inference. On the other hand, RoseTTAFold, with a limited number of predicted structures, primarily differs due to the low confidence level (pLDDT score) in the predictions by the model itself, making it unable to distinguish between the two types of structures effectively. Within the AlphaFold, we observed that the versions with and without MSA distributions are highly similar, and the structural similarity without MSA input is even higher than with MSA input. In other words, although there is no significant difference in MSA input, it still provides some discrimination in the model’s input.

**Figure 6:**
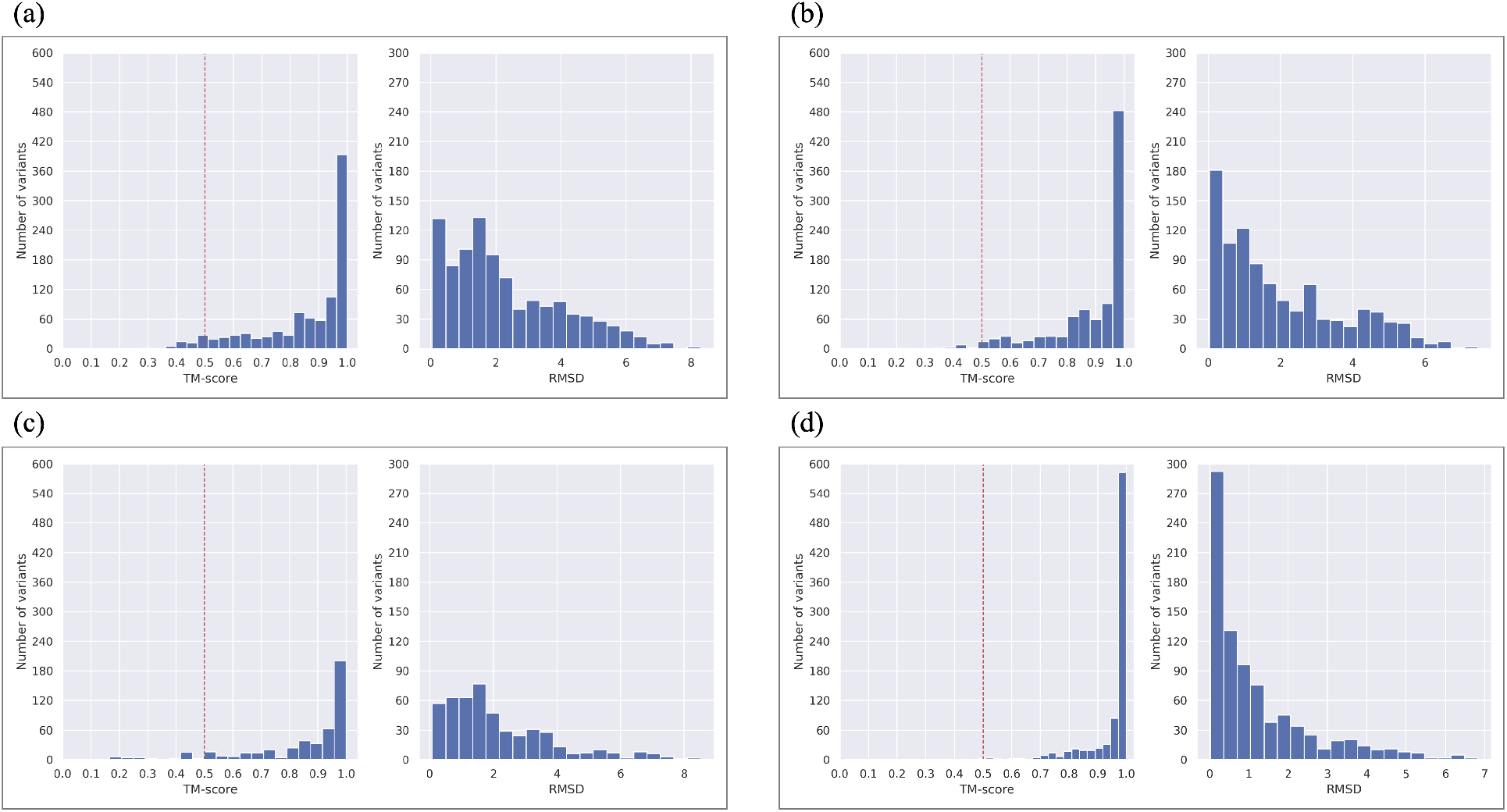
TM-align results between reference structures and alternative structures. (a) AlphaFold (b) AlphaFold without MSA (c) RoseTTAFold (d) ESMFold

By comparing the pLDDT score of alternative structures with their corresponding reference structures, we observed a high correlation between them (Figure 7(a)). Therefore, we selected the pLDDT score of the reference structure to compare with the TM-score. When a model has a high confidence level in its predictions, it becomes evident that the reference and variant sequences are more similar. However, we found 26 predicted structures where the model exhibits a certain degree of confidence (pLDDT > 70), but the reference structure and alternative structure significantly differ (TM-score < 0.5) (Figure 7(b)). We first ensure that these reference structures exhibit a high degree of similarity compared to the experimental structures. Excluding the two structures with TM-score < 0.7, the remaining structures are considered to have changes most likely related to nsSNV. Among them, MLH1 and its 19 variants are the most prominent instances.

**Figure 7:**
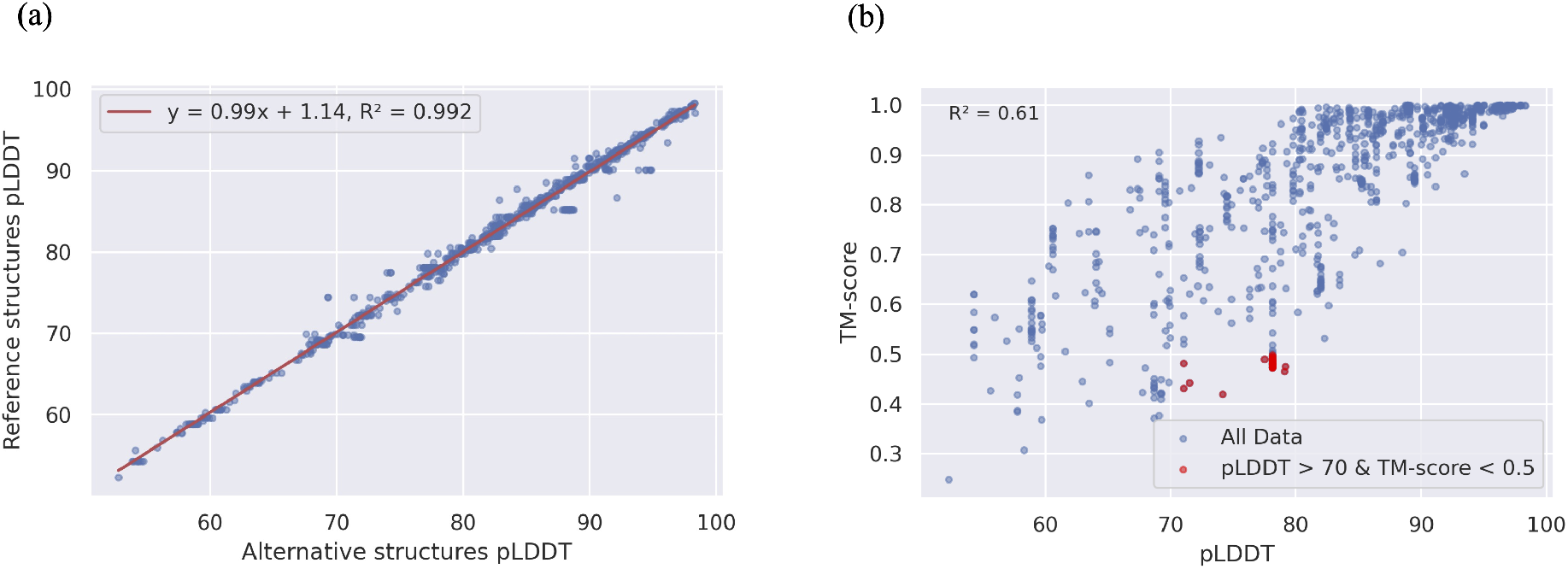
Relation between pLDDT score and TM-score. (a) High correlation for pLDDT score between reference structures and alternative structures. (b) Scatter plot for reference pLDDT score and its corresponding TM-score between predicted structures of reference sequences and alternative sequences. The red dots represent the cases we believe will most likely be distinctive for the structure prediction tool.

After comparing the structures in visualization tools, we found that the main differences in their structures arise from the regions of disordered regions. As shown in Figure 8, MLH1 contains two major foldable domains, and one aligns well with the experimental structure. After the disordered region (position 355-378), although the folding remains similar, the protein structures are spatially too distant to be aligned successfully, resulting in a lower TM-score. Additionally, we observed that the variation from alanine to glutamic acid at position 22 does not impact the structure prediction.

**Figure 8:**
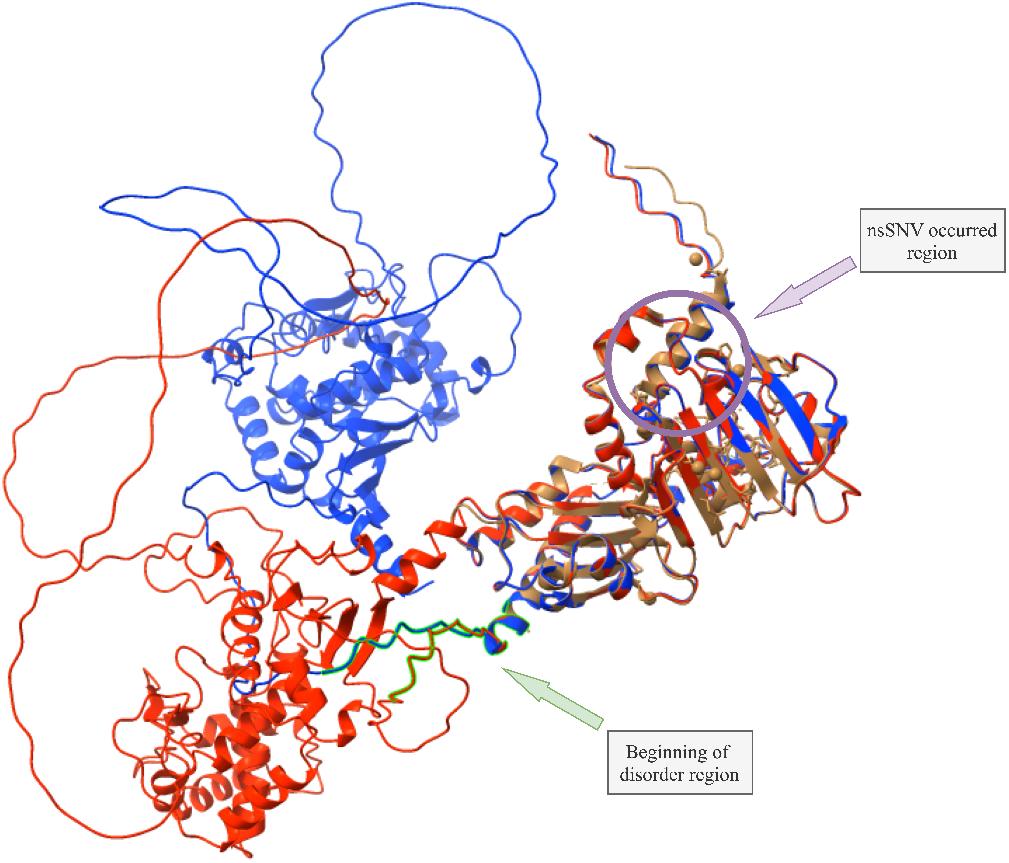
Visualization of MLH1. The brown structure is the best experimental structure after performing ClustalW. The blue structure is the predicted structure of the reference sequence, and the red one is the predicted structure of the alternative (A21E) sequence. All structures are aligned to the experimental structure. The visualization is using UCSF ChimeraX [25].

### 3.4 AlphaMissense

After the release of AlphaMissense, we included it in evaluating the selected data. In the predictions from AlphaMissense, 842 were classified as pathogenic, 69 as ambiguous, 48 as benign, and five cannot be mapped to the database among the selected variants. Although the model architecture is similar to AlphaFold, its pathogenicity assessment does not directly indicate structural changes. After all, we can see that the nsSNVs we chose are highly correlated with the AlphaMissense assessment, with 88% of them being identified as pathogenic variants by AlphaMissense. However, none of the three tools predicted structural changes in any selected variants in this study.

## 4 Conclusion

In this study, we compiled a dataset with a large number of pathogenic nsSNVs and executed structure prediction using three tools. We verified AlphaFold’s capability by choosing experimental structures that are most similar to the reference sequence utilizing ClustalW. Subsequently, we narrowed our discussion to a subset where the tools exhibited confidence and structural changes. AlphaFold reported low TM-align scores for some nsSNVs, more than the other tools. However, we found that in most cases, the inability to align structures between predicted structures of reference sequences and alternative sequences was not solely due to changes in amino acids but resulted from differentiation in the spatial orientation caused by disordered regions, leading to a decrease in TM-score. Allowing separate protein structure predictions for each domain might help avoid the effects of these disordered regions. In summary, the analyses conducted in this study indicated the current structure prediction tools cannot identify nsSNVs that might disrupt protein structure.

## References

[1] Brian Kuhlman and Philip Bradley. “Advances in protein structure prediction and design”. In: Nature Reviews Molecular Cell Biology 20.11 (2019), pp. 681–697. ISSN: 1471-0080. DOI: 10.1038/s41580-019-0163-x.

[2] D. Liebschner et al. “Macromolecular structure determination using X-rays, neutrons and electrons: recent developments in Phenix”. In: Acta Crystallogr D Struct Biol 75.Pt 10 (2019), pp. 861–877. ISSN: 2059-7983 (Electronic) 2059-7983 (Linking). DOI: 10.1107/S2059798319011471.

[3] Andriy Kryshtafovych et al. “Critical assessment of methods of protein structure prediction (CASP)—Round XIV”. In: Proteins: Structure, Function, and Bioinformatics 89.12 (2021), pp. 1607–1617. ISSN: 0887-3585. DOI: 10.1002/prot.26237.

[4] John Jumper et al. “Highly accurate protein structure prediction with AlphaFold”. In: Nature 596.7873 (2021), pp. 583–589. ISSN: 1476-4687. DOI: 10.1038/s41586-021-03819-2.

[5] Minkyung Baek et al. “Accurate prediction of protein structures and interactions using a three-track neural network”. In: Science 373.6557 (2021), pp. 871–876. DOI: doi:10.1126/science.abj8754.

[6] Zeming Lin et al. “Evolutionary-scale prediction of atomic-level protein structure with a language model”. In: Science 379.6637 (2023), pp. 1123–1130. DOI: doi:10.1126/science.ade2574.

[7] M. S. Hassan et al. “Evaluation of computational techniques for predicting non-synonymous single nucleotide variants pathogenicity”. In: Genomics 111.4 (2019), pp. 869–882. ISSN: 1089-8646 (Electronic) 0888-7543 (Linking). DOI: 10.1016/j.ygeno.2018.05.013.

[8] S. Iqbal et al. “Comprehensive characterization of amino acid positions in protein structures reveals molecular effect of missense variants”. In: Proc Natl Acad Sci U S A 117.45 (2020), pp. 28201–28211. ISSN: 0027-8424 (Print) 0027-8424. DOI: 10.1073/pnas.2002660117.

[9] Lukas Gerasimavicius, Benjamin J. Livesey, and Joseph A. Marsh. “Loss-of-function, gain-of-function and dominant-negative mutations have profoundly different effects on protein structure”. In: Nature Communications 13.1 (2022), p. 3895. ISSN: 2041-1723. DOI: 10.1038/s41467-022-31686-6.

[10] M. A. Pak et al. “Using AlphaFold to predict the impact of single mutations on protein stability and function”. In: PLoS One 18.3 (2023), e0282689. ISSN: 1932-6203. DOI: 10.1371/journal.pone.0282689.

[11] S. Ittisoponpisan et al. “Can Predicted Protein 3D Structures Provide Reliable Insights into whether Missense Variants Are Disease Associated?” In: J Mol Biol 431.11 (2019), pp. 2197–2212. ISSN: 0022-2836 (Print) 0022-2836. DOI: 10.1016/j.jmb.2019.04.009.

[12] H. Keskin Karakoyun et al. “Evaluation of AlphaFold structure-based protein stability prediction on missense variations in cancer”. In: Front Genet 14 (2023), p. 1052383. ISSN: 1664-8021 (Print) 1664-8021. DOI: 10.3389/fgene.2023.1052383.

[13] Anastassis Perrakis and Titia K Sixma. “AI revolutions in biology”. In: EMBO reports 22.11 (2021), e54046. ISSN: 1469-221X. DOI: 10.15252/embr.202154046.

[14] Melissa J Landrum et al. “ClinVar: improving access to variant interpretations and supporting evidence”. In: Nucleic Acids Research 46.D1 (2017), pp. D1062–D1067. ISSN: 0305-1048. DOI: 10.1093/nar/gkx1153.

[15] Kai Wang, Mingyao Li, and Hakon Hakonarson. “ANNOVAR: functional annotation of genetic variants from high-throughput sequencing data”. In: Nucleic Acids Research 38.16 (2010), e164–e164. ISSN: 0305-1048. DOI: 10.1093/nar/gkq603.

[16] P. C. Ng and S. Henikoff. “SIFT: Predicting amino acid changes that affect protein function”. In: Nucleic Acids Res 31.13 (2003), pp. 3812–4. ISSN: 0305-1048 (Print) 0305-1048. DOI: 10.1093/nar/gkg509.

[17] I. A. Adzhubei et al. “A method and server for predicting damaging missense mutations”. In: Nat Methods 7.4 (2010), pp. 248–9. ISSN: 1548-7091 (Print) 1548-7091. DOI: 10.1038/nmeth0410-248.

[18] Eugene V. Davydov et al. “Identifying a High Fraction of the Human Genome to be under Selective Constraint Using GERP++”. In: PLOS Computational Biology 6.12 (2010), e1001025. DOI: 10.1371/journal.pcbi.1001025.

[19] Rolf Apweiler et al. “UniProt: the Universal Protein knowledgebase”. In: Nucleic acids research 32.Database issue (2004), pp. D115–D119. ISSN: 1362-4962 0305-1048. DOI: 10.1093/nar/gkh131.

[20] H. M. Berman et al. “The Protein Data Bank”. In: Nucleic Acids Res 28.1 (2000), pp. 235–42. ISSN: 0305-1048 (Print) 0305-1048. DOI: 10.1093/nar/28.1.235.

[21] J. D. Thompson, D. G. Higgins, and T. J. Gibson. “CLUSTAL W: improving the sensitivity of progressive multiple sequence alignment through sequence weighting, position-specific gap penalties and weight matrix choice”. In: Nucleic Acids Res 22.22 (1994), pp. 4673–80. ISSN: 0305-1048 (Print) 0305-1048. DOI: 10.1093/nar/22.22.4673.

[22] Yang Zhang and Jeffrey Skolnick. “TM-align: a protein structure alignment algorithm based on the TM-score”. In: Nucleic Acids Research 33.7 (2005), pp. 2302–2309. ISSN: 0305-1048. DOI: 10.1093/nar/gki524.

[23] Y. Zhang and J. Skolnick. “Scoring function for automated assessment of protein structure template quality”. In: Proteins 57.4 (2004), pp. 702–10. ISSN: 0887-3585. DOI: 10.1002/prot.20264.

[24] O. Carugo and S. Pongor. “A normalized root-mean-square distance for comparing protein three-dimensional structures”. In: Protein Sci 10.7 (2001), pp. 1470–3. ISSN: 0961-8368 (Print) 0961-8368. DOI: 10.1110/ps.690101.

[25] E. F. Pettersen et al. “UCSF ChimeraX: Structure visualization for researchers, educators, and developers”. In: Protein Sci 30.1 (2021), pp. 70–82. ISSN: 0961-8368 (Print) 0961-8368. DOI: 10.1002/pro.3943.

